# An *in vivo* strategy for knockdown of circular RNAs

**DOI:** 10.1101/2020.01.23.916809

**Authors:** Nagarjuna Reddy Pamudurti, Ines Lucia Patop, Aishwarya Krishnamoorthy, Reut Ashwal-Fluss, Osnat Bartok, Sebastian Kadener

**Author notes:** To whom correspondence should be addressed Sebastian Kadener. These two authors contributed equally to this work (Nagarjuna Reddy Pamudurti, Ines Lucia Patop).

## Abstract

Exonic circular RNAs (circRNAs) are highly abundant and evolutionarily conserved RNAs generated mostly from exons of protein-coding genes. Assaying the functions of circRNAs is not straightforward as common approaches for circRNA depletion tend to also alter the levels of mRNAs generated from the hosting gene. Here we describe a methodology for specific knockdown of circRNAs *in vivo* with tissue and cell resolution. We also describe an experimental and computational platform for determining the potential off-target effects as well as for verifying the obtained phenotypes. Briefly, we utilize miRNA-derived shRNAs targeted to the circRNA-specific back-splice junction to specifically downregulate the circRNA. We utilized this methodology to downregulate five circRNAs that are highly expressed in the fly nervous system. There were no effects on levels of the linear RNA or any RNA with complementarity to the expressed shRNA. Interestingly, downregulation of circCtrip resulted in developmental lethality that was recapitulated with a second shRNA. Moreover, we found that downregulation of individual circRNAs caused specific changes in the fly head transcriptome, suggesting roles for these circRNAs in the fly nervous system. Together, our results provide a methodological approach that enables the comprehensive study of circRNAs at the organismal and cell levels.

## INTRODUCTION

Exonic circular RNAs (circRNAs) are a highly abundant type of RNA produced through circularization of specific exons in a process known as back-splicing ^1–4^. circRNAs are expressed in tissue- and development stage-specific ways, independently of the expression of the hosting gene ^5^. Indeed, abundant circRNAs can control expression of their hosting genes in *cis* by deviating transcriptions towards circRNA production ^6^. circRNAs are produced by the spliceosome and their production is driven by inverted repeat sequences in the RNA or RNA binding proteins ^6–12^. In addition, some circRNAs also produce proteins ^13–15^. The circRNA-encoded peptides usually share start codons with their hosting genes and might be important in synapse and muscle functions ^13^. circRNAs are particularly enriched in neural tissue ^16–19^. Moreover, circRNA levels increase with age in the brains of mice and flies as well as in worms ^18, 20, 21^ and are affected by neuronal activity^19^. These observations suggest important roles for circRNAs in the brain. For example, mice depleted of the circRNA CDR1as have abnormal gene expression in the brain and specific behavioral defects ^22^. Recent work has identified a handful of circRNAs that function in *trans*: The circRNAs derived from CDR1as and *sry* likely regulate miRNA function and/or localization ^22–25^. Other circRNAs titrate or transport proteins and might be important for cancer development ^6, 26, 27^. circRNAs can also mediate responses to viral infections ^28–31^

Despite steady advances, the circRNA field faces one main obstacle: It is usually not possible to downregulate the amount of a given circRNA without altering the levels of the linear RNA produced from the same hosting gene. Also, as circRNA production can compete with linear RNA splicing, it is difficult to separate the potential *cis* and *trans* functions of the circRNA. Recently, the Rajewsky lab generated the first animal (mouse) without a single circRNA (CDR1as). They did so by deleting the locus from which circRNA is generated. This was possible because the CDR1as locus is unusual as it does not encode a linear RNA. In cell culture studies, shRNAs (mostly transiently transfected) can alter circRNA levels, but shRNA-mediated knockdown is often inefficient (especially in mammalian systems) and can result in undesired silencing of the linear mRNA transcript encoded by the same locus or in other off-target effects. Deletion of intronic sequences responsible for exon circularization have also been used ^32^, this approach is laborious and would be difficult (or impossible) to use to globally screen for functions of circRNAs and it does not allow for tissue specific resolution. In addition, one cannot differentiate between *cis* and *trans* functions of circRNAs using this technique, and it can probably only be used for circRNAs that act only in *trans.* This is because impairing the biogenesis of circRNAs whose production regulates the expression of the hosting gene will result by definition in changes on the levels of the linear mRNA.

Here we adapted the use of shRNAs in order to specifically target circRNAs with cell and tissue resolution in *Drosophila*. Our methodology takes advantage of the functional separation of the miRNA and shRNA systems in flies and targets miRNAs derived from shRNAs specifically to the back-splicing junction. This allows us to achieve specific knockdown without altering the levels of the mRNA generated from the host gene. We demonstrated the utility of this method by targeting five highly expressed circRNAs. To determine potential off-targets we generated and sequenced RNA-seq libraries from the heads of these and control strains. We did not detect effects on any non-targeted RNAs with perfect or seed-like complementarity to the shRNA. Interestingly, downregulation of one of the targeted circRNAs, circCtrip, resulted in developmental lethality that we recapitulated with a second shRNA. Moreover, we found that downregulation of individual circRNAs led to specific changes in the fly head transcriptome, suggesting specific roles for these particular circRNAs in regulation of gene expression. Together, our results provide a methodological approach that enables the comprehensive study of circRNAs at the organismal and cell levels.

## RESULTS

### circRNAS can be specifically downregulated by miRNA-derived shRNAs *in vivo*

To knockdown circRNAs *in vivo* we generated flies that express shRNAs directed against individual circRNA-specific back-splicing junctions (Fig. 1A). For shRNA expression, we utilized a vector based on a miRNA-like precursor (miR-1 ^36^). For these experiments, we chose to target five circRNAs with high levels of expression in fly heads based on previous work ^6^. In all cases the shRNAs are expressed under the control of the GAL4/UAS system that allows temporal and spatial control of expression. We generated flies expressing the transgene under the control of a constitutive driver (*actin*-*Gal4*). In four of the five cases, we obtained viable flies (Supplementary Figure 1). We then evaluated the efficiency of the circRNA knockdown in fly heads and observed a specific and strong reduction in levels of the targeted circRNAs in fly heads (Figures 1B and C). For the four viable strains, expression of the shRNA led to more than 75% reduction in the level of the targeted circRNA (Figure 1B) without a noticeable effect on the levels of the other circRNAs (Figure 1C).

**Figure 1.**
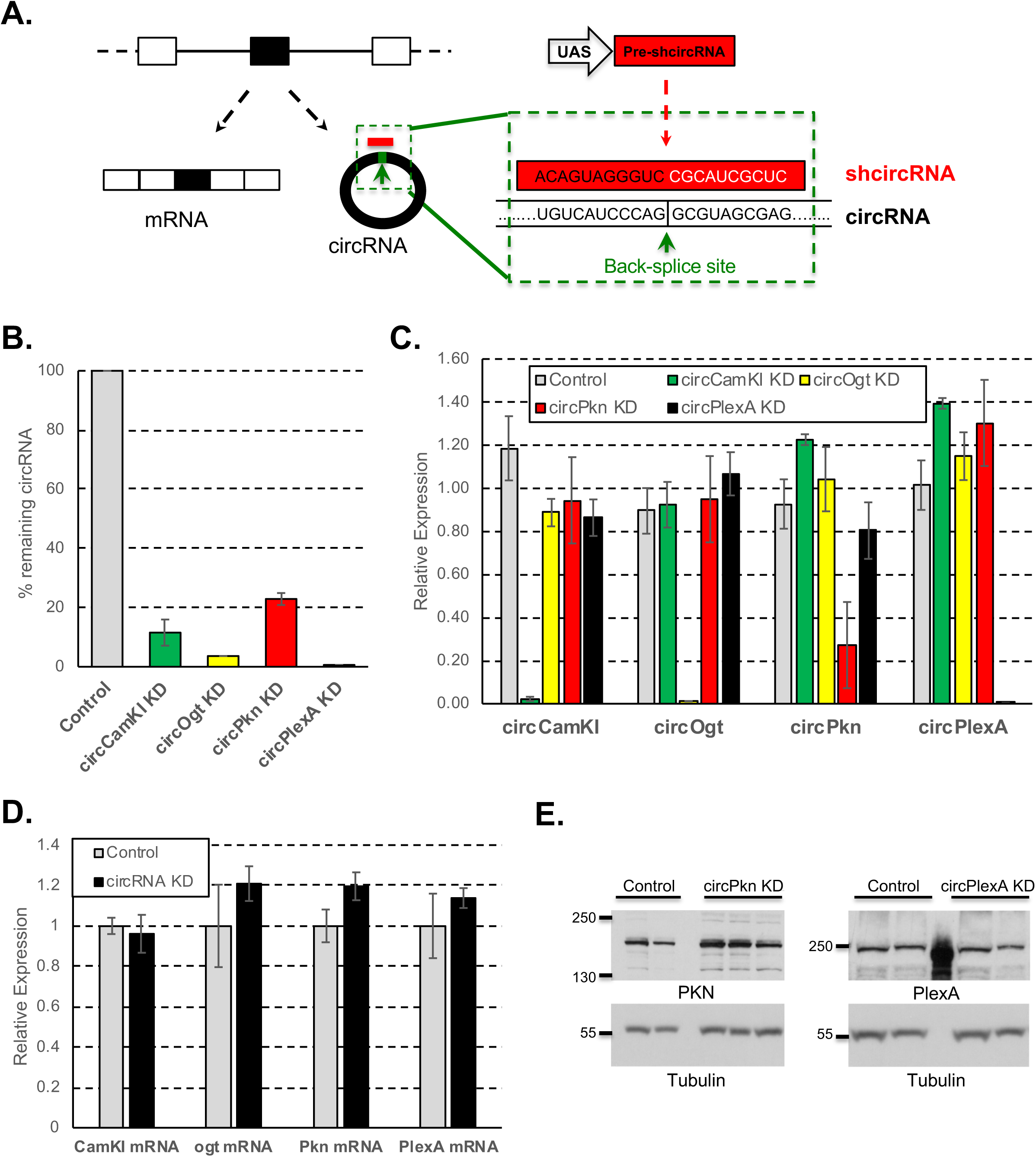
circRNAs can be specifically downregulated by genetically encoded shRNA *in vivo*. **A.** Scheme of the shRNA strategy to knockdown circRNAs *in vivo*. ShRNAs are generated from a UAS-based miR-1 like precursor (see methods). **B.** Relative levels of the targeted circRNAs in fly heads expressed as percentage of control (*actin-*Gal4 flies). Levels of the indicated circRNAs were assess by qRT-PCR using the *rp49* mRNA as normalization control (n=3, error bars represent standard error of the mean). **C.** Levels of the indicated circRNAs in the four knockdown strains. In all cases we utilized fly heads as source of material and assessed the levels of the circRNAs by qRT-PCR using *rp49* as normalization control (n=3, error bars represent standard error of the mean). **D.** RNAseq analysis of the indicated mRNAs in the indicated strains. In all cases we utilized *actin-*Gal4 flies as control (n=3, error bars represent standard error of the mean). **E.** Levels of the PKN and PLEXA proteins (and TUBULIN as loading control) as assayed by western blot in heads of control, circPkn and circPlexA KD strains.

### Most circRNA knockdown lines do not display identifiable off-targets effects

To further determine the specificity of the knockdowns, we generated and sequenced 3’-end libraries from heads of the four knockdown strains that did not cause embryonic lethality. The 3’-RNA-seq technique is highly reliable for determining mRNA levels of low abundance mRNAs ^34^. We observed that for all the assayed strains, the shRNAs did not significantly alter the levels of the mRNAs produced from the same locus (Figure 1D; Table S2).

Despite the lack of effect of the shRNAs on the level of the linear mRNAs, it is possible that translation of the hosting mRNAs could be impacted. We tested this possibility by analysis of two strains for which there are antibodies available against the proteins encoded by mRNAs generated from the hosting genes. Expression of the shRNAs targeting the back-splice junctions of circPkn and circPlexA strongly downregulated the targeted circRNAs without detectable effect on the levels of PKN or PLEXA proteins (Figure 1E). These experiments demonstrate that specifically targeting circRNA junctions with shRNA depletes the circRNA without any impact on hosting gene expression.

In addition, shRNAs could have non-specific effects on other mRNAs. To evaluate this possibility, we determined whether downregulated mRNAs in shRNA-expressing strains were enriched for sequence complementary to the seed of the shRNA or shRNA* utilizing the SYLAMER algorithm^35^. In none of our lines the downregulated mRNAs were enriched for any relevant seed sequences (Fig. 2A).

**Figure 2.**
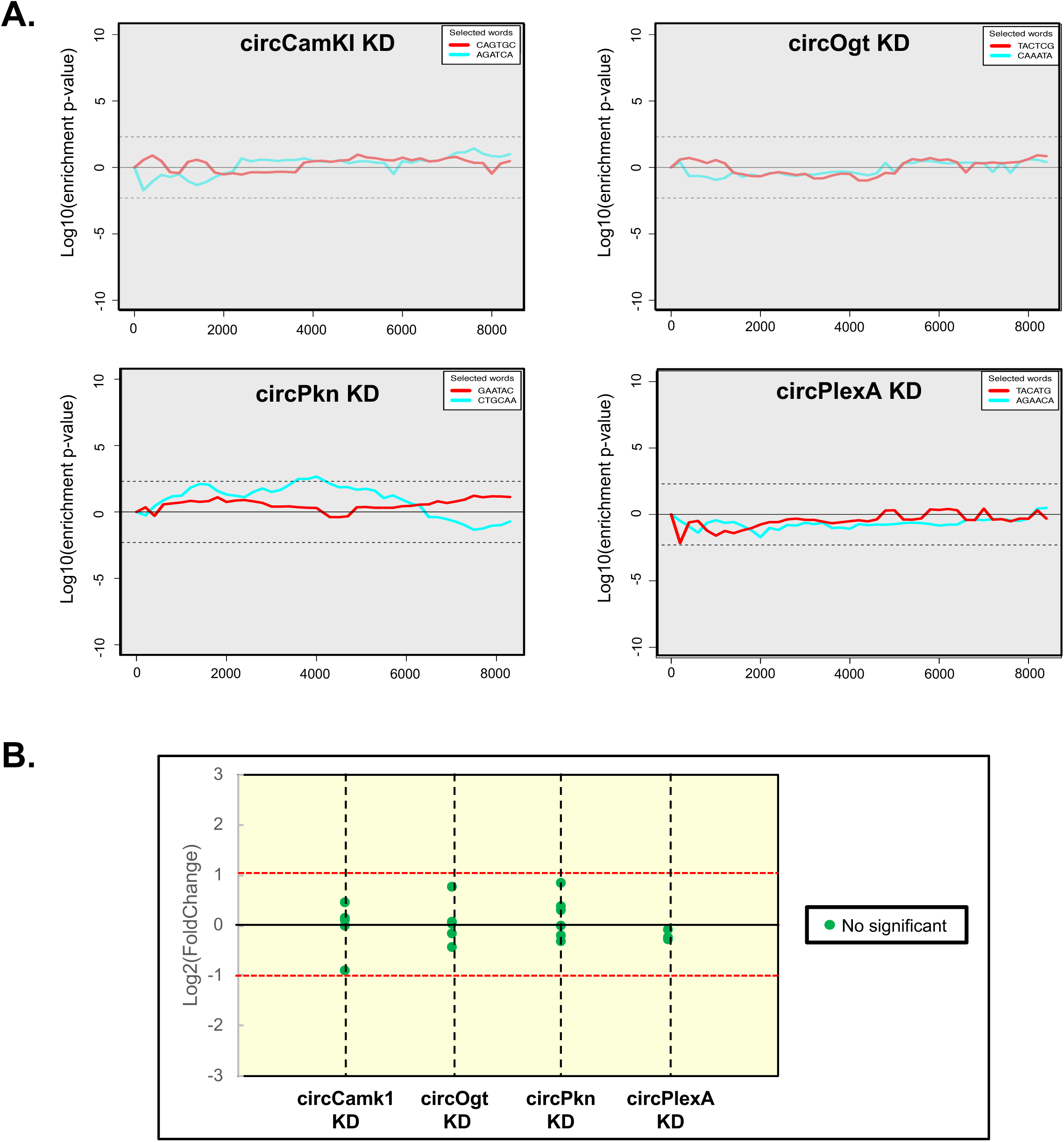
shRNAs targeting circRNAs have no detectable off targets. **A.** Assessment of off-targets of the indicated shRNAs by SYLAMER. Seed enrichment was calculated for the genes differentially expressed upon downregulation of circCamkI, circOgt, circPkn and circPlexA. shRNA and shRNA* seed sequences shown in red and blue, respectively. Genes were sorted from downregulated to upregulated. For all knockdowns we did not observed any shRNA seed enriched among the downregulated genes. **B.** Expression of putative off-targets genes in circRNA KD lines. Potential off-targets genes were selected using blast of the shRNA sequence against the *Drosophila* transcriptome. 3’ RNA-seq data was used to determine the expression level of each putative off-targets gene relative to control line (*actin*-Gal4) and presented as log2(Fold change). None of the mRNAs displayed statistically significant differences (fold change >1.5 and corrected pvalue<0.05).

It is also possible that shRNAs could target mRNAs with more extensive base complementarity in any part of the mRNA. To test this, we identified mRNAs with 12 or more bases complementary to the shRNAs and determined whether any were downregulated upon expression of shRNA. To do so we created a script that automatically blast the shRNA sequence into NCBI and output the corresponding matches (see methods section). Indeed, none of the mRNA with base complementarity to the shRNA display significant changes in expression upon silencing of the circRNA (Fig. 2B and Table S3). Together, these results demonstrate that these shRNAs act specifically and do not display observable off-target effects.

### circRNA knockdown provokes specific changes in the head transcriptome

We then analyzed more carefully the mRNAs differentially expressed in fly heads upon expression of the four non-lethal shRNAs. We found that downregulation of specific circRNAs provoked specific changes in the head transcriptome (Fig. 3A and Table S2 and S4). Interestingly, for most of the circRNA knockdown strains, genes belonging to specific Gene Ontology (GO) terms were significantly enriched in the differentially expressed genes (Figure 3B and Table S5). For example, knockdown of circCamk1 provoked changes in genes related with sensory perception of smell and sugar metabolism. For the circOgt KD line, the enriched terms included GO terms related with signaling receptor activity, while for the circPkn KD line, we found GO terms related to muscle function and response to pheromone. The differentially expressed genes in flies in which circPlexA was knocked-down were enriched for GO terms related to immune response. Interestingly, we found some shared terms between the different fly strains. Among them were neurotransmitter binding and receptor regulator activity. Overall, the gene expression results suggest that these circRNAs might have different physiological functions.

**Figure 3.**
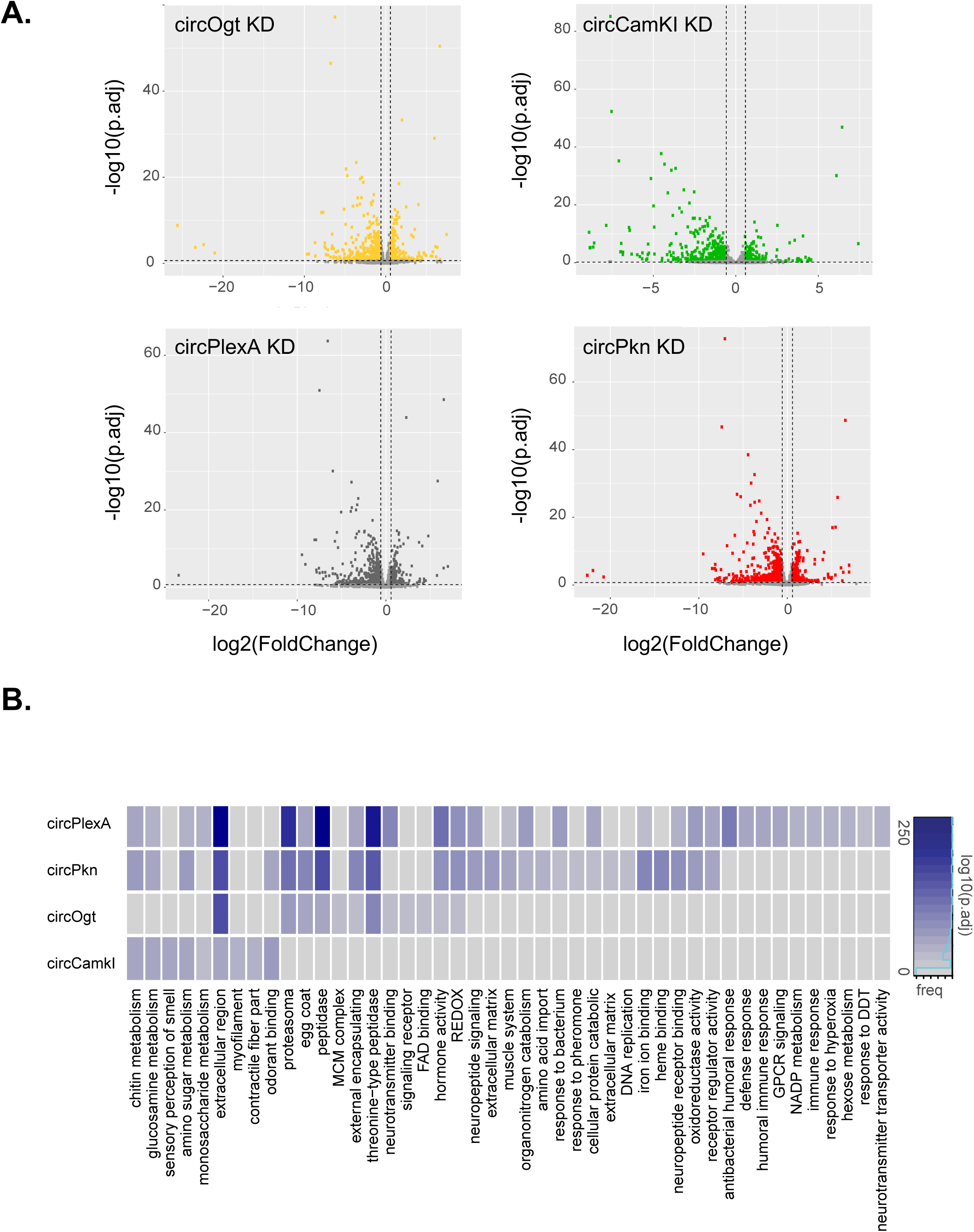
circRNA KD modulates specific set of gene expression. **A.** Differentially expressed genes in different circRNA KD flies in comparison to a control flies (*actin*-Gal4; n=3 for each strain). Each plotted point represents the differential expression results for an individual gene. The x axis shows the log2 of fold change between the knockdown and the control strain fly. The y axis shows the –log10(adjusted p-value) for the fold change per gene. **B.** Gene Ontology (GO) terms significantly enriched (FDR < 0.1) among genes differentially expressed between the heads of control and circRNA-KD flies. Color represent -log10 of adjusted p-value. Redundant terms were curated manually and names were simplified for clarity reason. Complete results are in Table S5.

### Knockdown of circCtrip leads to developmental phenotypes

Expression of an shRNA designed to deplete circCtrip led to very strong developmental lethality (Supplementary Figure 1). Moreover, the few flies that did eclose from this strain (*actin-Gal4*; UAS-shcircCtrip/CG14656) had locomotion problems and died within a few days (Video S1). We utilized a GFP-labeled balancer chromosome to determine the timing of the lethality. We found that expression of the shRNA against circCtrip caused lethality at the pupal stage (Fig. 4A). Consistent with this, analysis of previously published gene expression data of flies over the course of development (Westholm et al., 2014) showed that circCtrip is expressed at the larval and pupal stages (Fig. 4B).

**Figure 4.**
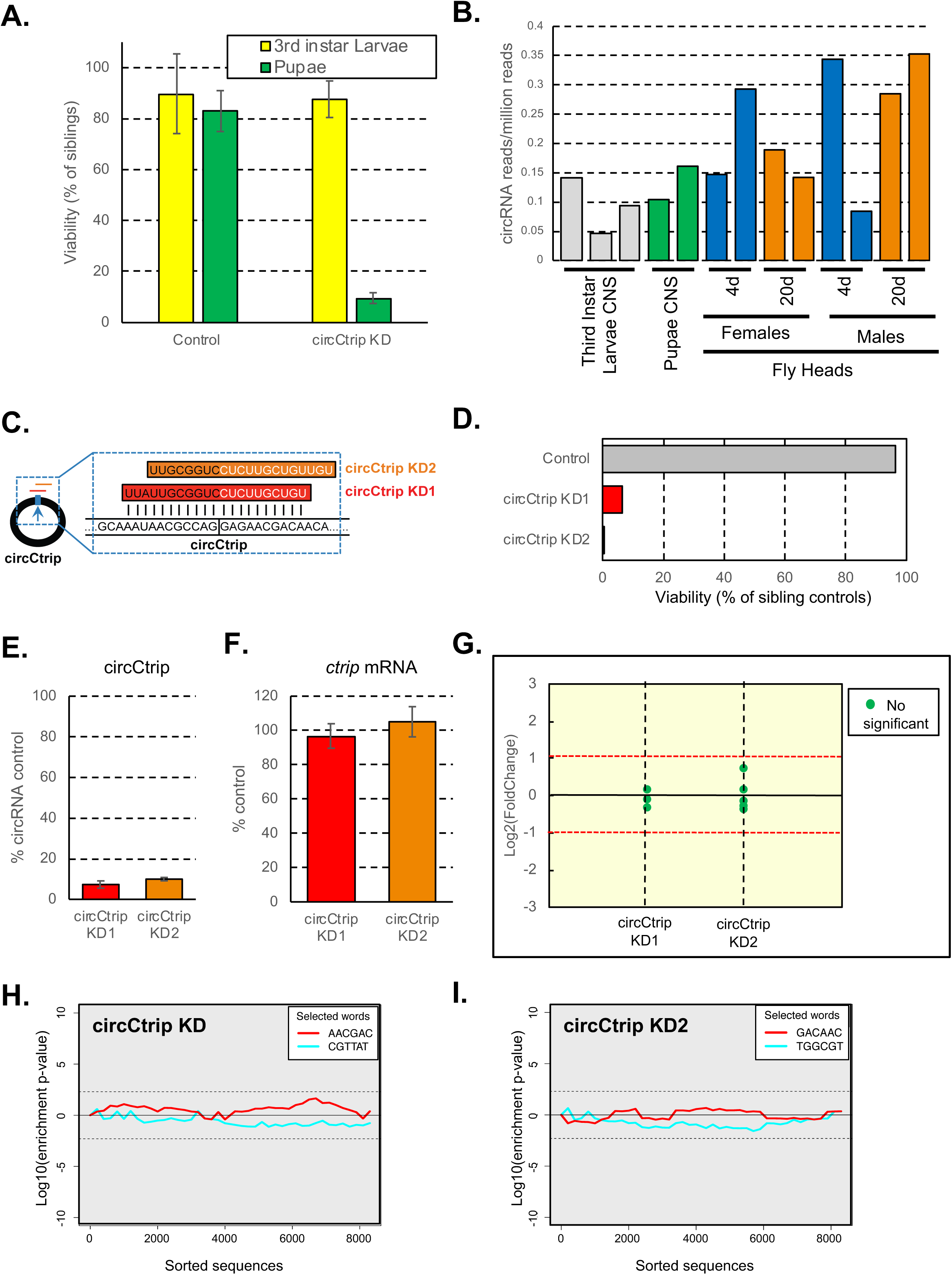
circCtrip is essential for proper development. **A.** Viability of circCtrip knockdown compared to their sibling controls. In green, percentage of pupae and in yellow 3rd instar larvae developed from sorted 1^st^ instar larvae. circCtrip KD flies show more than 90% of larval lethality when compared to the control flies (*actin*-Gal4). For each KD strain we normalized the data to the one of the siblings (GFP+) individuals (n = 3 sets of 50 embryos each). **B.** Expression of circCtrip at different developmental stages. Data was extracted from (23) and reanalyzed for circRNA expression as described in (23) Y axis indicates number of circRNA reads per million reads. **C.** Schematic representation of the additional shRNAs (shRNA-2) designed against circCtrip. **D.** Developmental viability (% of siblings control) of flies expressing the second shRNAs against circCtrip. **E.** qRT-PCR evaluation of the efficiency of knockdown of the circRNA by the use of the different shRNA lines in the fly CNS. In all cases we utilized the *elav*-Gal4 driver to express two different shRNAs for circCtrip. We then determined the efficiency of the knockdown by comparing to the levels of the assayed circRNA in the control strain (*elav*-Gal4). Data was normalized to 18S rRNA and *rp49* (n=3, error bars represents standard error of the mean). **F.** 3’ RNA-seq data was used to determine the expression level of Ctrip mRNA in circCtrip KD lines (n=3, error bars represents standard error of the mean). **G.** Expression of putative off-targets genes in both circCtrip KD lines. Potential off-targets genes were selected using blast of the shRNA sequence against the drosophila transcriptome. 3’ RNA-seq data was used to determine the expression level of each putative off-targets gene relative to control line (*elav*-Gal4) and presented as log2(Fold change). The different colors of the dots represent the statistical significance of the differences. **H and I,** assessment of off-targets of the indicated shRNAs by SYLAMER. Seed enrichment was calculated for the genes differentially expressed upon downregulation of circCtrip using each shRNA. shRNA and shRNA* seed sequences shown in red and blue, respectively. Genes were sorted from downregulated to upregulated. For both knockdowns we did not observed any shRNA seed enriched among the downregulated genes.

To rule out off target effects, we generated a new fly line expressing a second shRNA targeting circCtrip. The new shRNA was designed to be perfectly complementary to the circRNA, but the target site was shifted and hence potential off target effects should be reduced (Figure 4C). We found that the second shRNA construct targeting circCtrip also resulted in developmental lethality (Figure 4D). To confirm that expression of these two shRNAs did indeed downregulate circCtrip, we generated flies in which the shRNAs were expressed specifically in the fly nervous system by using the *elav-Gal4* driver. We observed a strong downregulation of circCtrip in both shRNA-expressing lines (Figure 4E).

To completely rule out the possibility that the developmental lethality of the circCtrip shRNAs is due to off-target effects, we prepared and sequenced 3’-RNA-seq libraries from heads of the circCtrip and control strains. We did not detect significant effects of either shRNA on the levels of *ctrip* mRNA (Figure 4F). In addition, the genes with complementarity to these shRNAs were not differentially expressed upon the expression of either shRNA targeting circCtrip (Figure 4G; Table S6). Last but not least we tested using SYLAMER for potential miRNA-like effects of the shRNA or shRNA*. We indeed did not observe any enrichment for mRNAs containing the seed sequence among those downregulated upon expression of any of the two shRNAs against circCtrip (Figure 4H). These results demonstrate that that shRNA targeting circCtrip have no detectable off-target effects in fly heads and that this circRNA is necessary for proper development.

## DISCUSSION

In this manuscript, we adapted the use of genetically encoded shRNAs to downregulate specific circRNAs in *Drosophila*. To do so, we designed miRNA-derived shRNAs that target the circRNA-specific back-splicing junctions of five highly expressed circRNAs. Expression of these shRNAs led to very strong and specific downregulation of the targeted circRNAs with no effects on the levels of the linear RNA and in two cases protein generated from the same loci. In addition, we did not detect any changes in expression of genes with complementarity to the shRNA or with seed-like sequences in their 3’ UTRs, demonstrating that the utilized shRNAs have no detectable off-target effects. We utilized this approach to downregulate five highly expressed circRNAs *in vivo*. We found that downregulation of circCtrip resulted in developmental lethality that was recapitulated with a second shRNA. Moreover, we found that downregulation of individual circRNAs caused specific changes in gene expression. In sum, the methodology described here will allow investigation of the functions of circRNAs *in vivo*, and we have established quality standards for detection of specific and off-target effects of the shRNAs.

Although circRNAs are very abundant, especially in neural tissue, only a few studies have assessed the functionalities of these molecules *in vivo.* A main obstacle to functional experiments is the lack of tools for specifically manipulating circRNA levels. As expression of circRNAs generally depends on inverted RNA repeats or RNA binding proteins, it is theoretically possible to modulate circRNA levels by deleting those elements from the DNA (and hence RNA) by CRISPR. However, this type of manipulation might also alter the levels of the mRNA generated from the locus. Perturbation of the expression of the linear RNA might reflect a function in *cis* of the circRNA production (regulation of linear mRNA production) or an unwanted effect of the manipulation. This makes deletion experiments non-optimal, in particular for circRNAs with high production rates that can potentially have functions in *cis* and *in trans*.

shRNAs target circRNAs after production allowing the study of *trans*-like functions of these molecules. However, shRNA-mediated silencing has potential problems, chiefly the targeting of RNAs other than the desired circRNA. In particular, depending on sequence, shRNAs can target linear RNAs generated from the circRNA-hosting locus. Limited complementarity (10-11 bases) can in some cases be enough to induce slicing of an mRNA or can inhibit protein production if the shRNA acts as a miRNA. Although this is a problem in mammals, it is less likely to be an issue in *Drosophila* in which the miRNA and shRNA pathways are compartmentalized as the AGO and DICER proteins are different in the two pathways ^37^. Indeed, we did not observe miRNA-like downregulation of the hosting or other mRNAs when any of our shRNAs were expressed. In addition, we did not see any changes in the levels of mRNAs sharing >12 bases with the expressed shRNA. These results strongly suggest that the shRNAs used here have no off-target effects, although it is not possible to completely rule out this possibility. When implementing our shRNA-based approach, it will be necessary to perform off-target analysis such as those described here.

Additional and concurrent knockdown or knockout approaches could corroborate the phenotypes we observed upon circRNA knockdown. These approaches could include the generation of additional shRNA-expressing strains that target slightly different regions of the back-splice junctions, the use of other systems for degrading specific RNAs (e.g., Cas13b ^38^), or the generation of flies with the circRNA knocked down by a different method (e.g., by CRISPR deletion of inverted repeats of RNA binding protein sites). The method described here constitutes a straightforward method for knockdown of circRNAs in a specific tissue or the whole organism that can be used as a first step in analysis of circRNA function.

Importantly, the observed knockdown is strong (more than 5-fold and in most cases almost total) and can be restricted in time and space. Spatial resolution is easy to achieve in *Drosophila* by the use of different GAL4 drivers ^39^. Indeed, here we restricted the expression of the shRNAs to neuronal tissue. Several systems that allow for temporal expression exist in *Drosophila.* The most commonly used involves the reversible inhibition of GAL4-based transgene expression by a temperature sensitive (ts) GAL80 inhibitor ^39^. We do not believe that this strategy of GAL4-induction (and hence silencing) will work to silence circRNAs, as circRNA expression dramatically increases at higher (29°C) temperatures ^6^. However, other systems such as the Geneswitch should efficiently induce the expression of UAS-based shRNA in a temporally regulated fashion ^39^..

In sum, this manuscript describes a platform for silencing of circRNA expression that includes methods for determination of the accuracy and specificity of the shRNA reagents. We generated several *Drosophila* lines in which specific circRNAs were targeted for degradation using shRNAs. We observed that circCtrip is essential for proper fly development, whereas the other four circRNAs evaluated might regulate gene expression. Together, this works presents a new approach for assessing the functions of circRNAs *in vivo*.

## MATERIALS AND METHODS

### Fly strains

Wild type flies that we used in this study are from the *w^1118^* strain (Bloomington *Drosophila* Stock Center Indiana, USA). For constitutive knockdown of circRNAs we utilized the *Actin-*Gal4 driver (stock number 3953, Bloomington *Drosophila* Stock Center, Indiana, USA). For neural specific knockdowns we utilized an *elav*-Gal4; UAS *Dcr2* strain that was generated by using *elav*-Gal4 (stock number 458, Bloomington *Drosophila* Stock Center, Indiana, USA) and UAS-*Dcr2* flies. All crosses were performed and raised at 25 °C.

### Generation of shRNA lines

To generate circRNA KD flies we designed oligonucleotides with perfect 21-nucleotide complementary sequence to the circRNA junction, annealed them, and ligated in to the linearized Valium20 vector with EcoR1 and Nhe1 restriction enzymes. Colonies were screened by PCR and the plasmid was purified and sequenced from positive colonies. These plasmids were sent for injection to BestGene Inc (CA, USA). The presence of potential off-targets was verified before generating these strains by performing Blast against the fly genome and transcriptome. We did not observe any perfect complementary of 16 bases or more for any of the shRNAs.

### Real Time PCR analysis

Total RNA was extracted from adult fly heads using TRI Reagent (Sigma) and treated with DNase I (NEB) following the manufacturer’s protocol. cDNA was synthesized from this RNA (using iScript and random primers, Bio-Rad) and was utilized as a template for quantitative real-time PCR performed with the C1000 Thermal Cycler Bio-Rad. The PCR mixture contained Taq polymerase (SYBR green Bio-Rad). Cycling parameters were 95 °C for 3 min, followed by 40 cycles of 95 °C for 10 s, 55 °C for 10 s and 72 °C for 30 s. fluorescence intensities were plotted versus the number of cycles by using an algorithm provided by the manufacturer. Primer efficiency was determined for all primers described in this study and was taken into account for the relative expression calculation. The sequences of all the primers used in this assay are detailed in Table S1.

### Assessment of developmental lethality

Ten homozygous shcircRNA male flies were crossed with 10 virgin female *actin*-Gal4 flies and transferred to new bottles every 3 days. The F1 progeny was separated based on their genotype (indicated by the presence of the marker/balancer CyO) and the number of males and females flies counted. We performed this assessment for each bottle for 9 days or until the totality of the F1 eclose.

### RNA libraries preparation for RNA-seq analysis

Total RNA was extracted using Trizol reagent (Sigma) and treated with DNase I (NEB) following the manufacturer’s protocol. Then, 0.5µg of total RNA was fragmented in FastAP buffer (Thermo Scientific) for 3min at 94^0^C, then dephosphorylated with FastAP, cleaned (using SPRI beads, Agencourt) and ligated to a linker1

(5Phos/AXXXXXXXXAGATCGGAAGAGCGTCGTGTAG/3ddC/, where XXXXXXXX is an internal barcode specific for each sample), using T4 RNA ligase I (NEB). Ligated RNA was cleaned-up with Silane beads (Dynabeads MyOne, Life Technologies) and pooled into a single tube. This mix then polyA+ selected (using Oligo(dT) beads, Invitrogen), RT was then performed for the pooled sample, with a specific primer (5’-CCTACACGACGCTCTTCC-3’) using AffinityScript Multiple Temperature cDNA Synthesis Kit (Agilent Technologies). Then, RNA-DNA hybrids were degraded by incubating the RT mixture with 10% 1M NaOH (e.g. 2ul to 20ul of RT mixture) at 70^0^C for 12 minutes. pH was then normalized by addition of corresponding amount of 0.5M AcOH (e.g. 4ul for 22 ul of NaOH+RT mixture). The reaction mixture was cleaned up using Silane beads and second ligation was performed, where 3’end of cDNA was ligated to linker2 (5Phos/AGATCGGAAGAGCACACGTCTG/3ddC/) using T4 RNA ligase I. The sequences of linker1 and linker2 are partially complementary to the standard Illumina read1 and read2/barcode adapters, respectively. Reaction Mixture was cleaned up (Silane beads) and PCR enrichment was set up using enrichment primers 1 and 2:

(5’AATGATACGGCGACCACCGAGATCTACACTCTTTCCCTACACGACGCTCTTCCGA TCT-3’, 5’-CAAGCAGAAGACGGCATACGAGATXXXXXXXXGTGACTGGAGTTCAGAC GTGTGCTCTTCCGATCT-3’, where XXXXXXX is barcode sequence) and Phusion HF MasterMix (NEB). 12 cycles of enrichment were performed. Libraries were cleaned with 0.7X volume of SPRI beads. Libraries were characterized by Tapestation. RNA was sequenced as paired-end samples, in a NextSeq 500 sequencer (Illumina).

### Western blot analysis

Fly heads (20 heads per sample) were collected on dry ice. Heads were homogenized in RIPA lysis buffer (50 mM Tris-HCl at pH 7.4, 150 mM NaCl, **1** mM EDTA, 1% NP-40 0.5% Sodium deoxycholate, and 0.1% sodium dodecyl sulfate (SDS), 1 mM DTT, supplemented by protease inhibitor cocktail and phosphatase inhibitors) using a motorized pestle. Head lysates were then centrifuged at maximum speed for 10 minutes and the supernatant was saved. lysates were boiled with protein sample buffer (Bio-Rad) and resolved by Criterion XT Bis-Tris gels (Bio-Rad). Antibodies used for western blotting: srabbit anti PKN was kindely provided by Prof. Rui Goncalo Martinho, (University of Algarve, Portugal, 1:1000), rabbit anti PlexA was kindly provided by Prof. Liqun Luo (Howard Hughes medical institute, Stanford university, 1:1000), mouse anti-tubulin (DM1A; SIGMA, 1:30,000).

### Determination of developmental stage at which circRNA KD provokes lethality

We used *actin*-Gal4/CyO-GFP to achieve ubiquitous knockdown of circRNAs and score out non-Gal4 expressing siblings by excluding GFP-expressing larvae. Virgin females of Gal4 and males of UAS-shRNA were crossed a day before. On the day of experiment, we transferred the crossed flies to embryo collection chamber with yeast paste on sucrose agar plates. Three sets of embryos (∼100 each) were collected, aligned in a straight line against a coverslip on a sucrose agar plate. Thus, aligned embryos are allowed to develop and hatch into larvae. We then identified, counted and separated based on the presence of GFP the 1^st^ instar larvae. Then, we counted the number of 3^rd^ instar larvae and the number of larvae that pupated. The data corresponding to the Non-Cyo (shcircRNA KD) was then normalized to the one of their siblings, averaged and plotted.

### Computational analysis

#### Gene expression analysis

RNA-seq reads were aligned to the genome and transcriptome (dm3) using tophat ^33^. Gene expression levels from 3’ DGE experiments were determined using ESAT tool ^34^ and differential expression analysis was performed with DEseq2. We considered genes with fold change>1.5 and pvalue<0.05 as significantly changing. *Actin*-Gal4 flies were used as a control for the lines expressing shRNA under actin promoter. In order to clean non-specific effects, we excluded from downstream analyses genes that change in all the circRNA KD strains. *elav*-Gal4; UAS-Dcr2 flies were used as a control for the lines expressing shRNA under *elav* promoter. clusterProfiler package was used for enrichment analysis of gene ontology. GO terms with p-value <0.1 (after FDR correction) were considered significant.

#### Determination of potential off-targets

We determined of targets using two different approaches. First, we use SYLAMER algorithm ^35^ to check for general off-target effect of the shRNA in the 3’ UTR regions of genes differentially regulated. To that end we sorted the genes 3UTR sequences by fold change (from downregulated to upregulated) and searched for enrichment of the miRNA seed of the shRNAs. Secondly, in order to obtain a list of general potential shRNA off target genes we blast all shRNA sequences against the *Drosophila* transcriptome. To this aim we created a python script using *qblast* from biopython to invokes the NCBI BLAST for each of the shRNAs sequences (https://github.com/ipatop/blast_for_shRNA_design). 3’ RNA-seq data was used to determine the expression level of each putative off-targets gene relative to control line.

## Acknowledgements

We thank the members of the Kadener lab for helpful discussions.

## Author Contribution

**NR** designed the shRNAs and performed most of the experiments; **IP** wrote the manuscript and performed the computational analysis; **AK** performed the crosses and the developmental experiments; **RA** generated the pipelines and conceptual framework for performing the off target assessments; **OB** performed the western blots and generated the RNAseq libraries; **SK** designed the experiments, supervised the project and wrote the manuscript.

## LEGEND OF SUPPLEMENTARY MATERIALS

**Figure S1.**
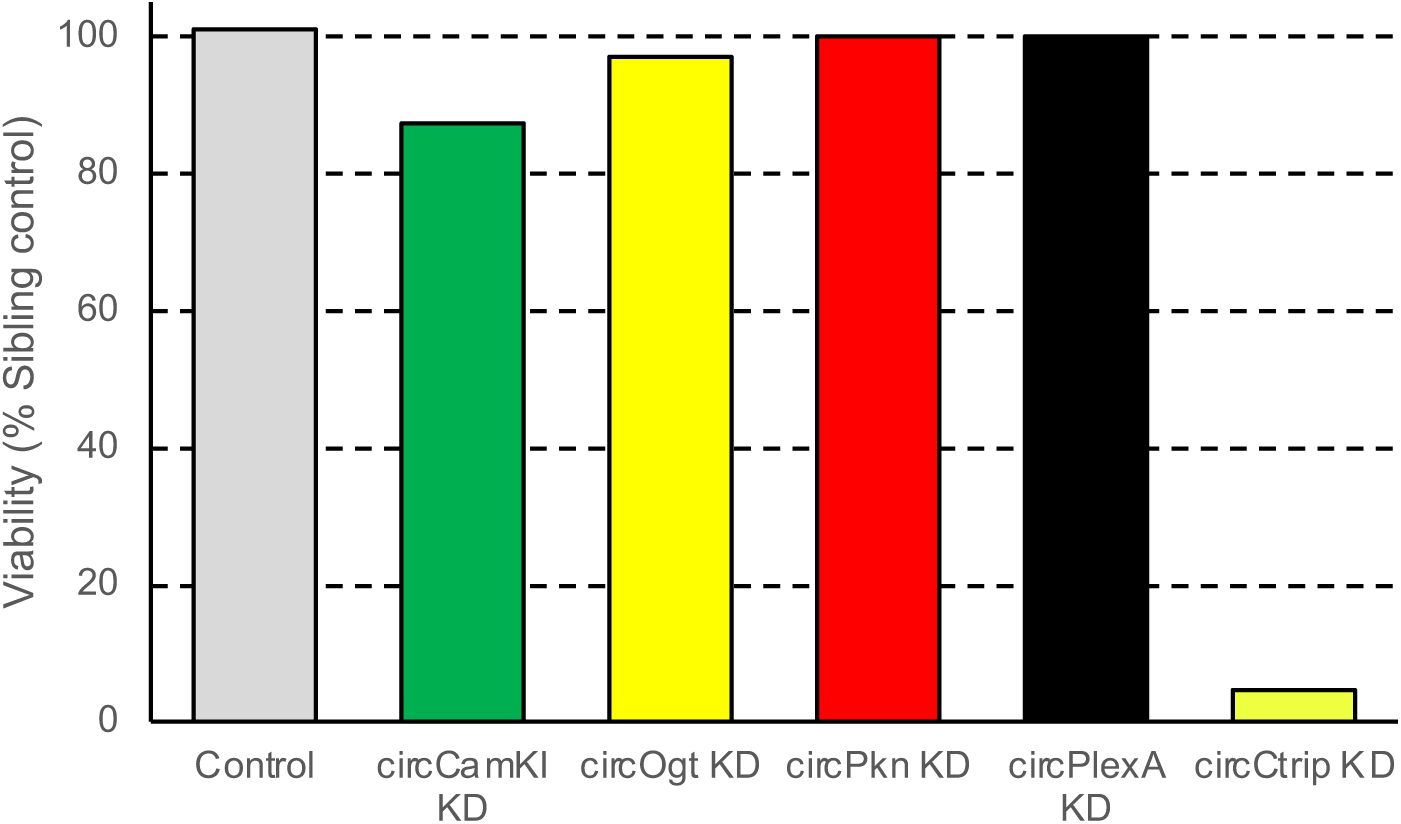
Viability of all the circRNA knockdown strain compared to their sibling controls. For each KD strain we normalized the data to the one of the Cyo individuals (In all cases the experiment was performed at least three times).

**Table S1. Oligonucleotides used in this study.**

**Table S2. Normalized gene expression for the circRNA knockdowns.** Results of the 3’-RNAseq for each circRNA KD. In all cases we utilize the *actin-gal4* driver to express the shRNA. The control is the *actin-*gal4 strain.

**Table S3. List of possible off-target genes and their expression changes in the different circRNA knockdown strains.** In all cases the displayed data was obtained by RNAseq and we note the log2(foldChange) and adjusted p-value for the respective knockdown. The data analyzed here is the same that is displayed in Table S2.

**Table S4. Genes differentially expressed upon knockdown of specific circRNAs.** Differential expression analysis of the 3’-RNAseq (Table S2) datasets. See methods for exact description of the analysis.

**Table S5. Gene Ontology (GO) term analysis of the differentially expressed genes upon specific circRNA knockdowns.** The displayed value for each GO term is -log10(pvalue adjusted) for each shRNA genotype

**Table S6. List of possible off-target genes and their expression changes in the two circCtrip KD strains.** In all cases the displayed data was obtained by RNAseq and we note the log2(foldChange) and adjusted p-value for the respective knockdown. The shRNAs directed against circCtrip were expressed under the control of the *elav-*gal4 driver. The control strain for this analysis was *elav-*gal4.

**Video S1. circCtrip KD escapers have impaired locomotion than control flies.** The middle tube in the video contains circCtrip KD escapers flies which have serious locomotive defects.

